# SCUBE-1 in The Diagnosis and Follow-Up Of Subclinical Atherosclerosis in The Experimental Diabetes Mellitus and Obesity Model

**DOI:** 10.1101/2023.10.03.560790

**Authors:** Mehmet KÖK, Rahime ASLANKOÇ, Özlem ÖZMEN, Devrim DORA, Hamit Yasar ELLIDAG

## Abstract

**BACKGROUND:** Obesity and diabetes mellitus (DM) is a metabolic disease that is a global problem, the most crucial complication of which is atherosclerotic cardiovascular disease resulting from endothelial dysfunction and accompanying platelet hyperactivity. The presence of the SCUBE-1 protein has been observed within vascular endothelial cells and platelets, both recognized for their pivotal involvement in the arterial thrombosis mechanism. The objective of this research is to examine the utility of serum SCUBE-1 levels in diagnosing and monitoring subclinical atherosclerosis in experimentally induced DM and obese rats.

**METHODS:** The study comprised a cohort of 28 male Sprague‒Dawleyrats, which encompassed the following groups: the obese group subjected to a high-fat diet (HFD), the TII-DM group administered HFD in combination with a single dose of streptozocin (STZ), the TI-DM group treated solely with STZ, and the control group. Serum SCUBE1 was analyzed using the enzyme-linked immunosorbent assay method, and caspase-3 (Cas-3), interleukin-6 (IL-6), interferon-gamma (INF-gamma) and superoxide dismutase (SOD) expression in the liver and pancreas of rats were evaluated using immunohistochemical methods.

**RESULTS:** Serum SCUBE1 levels were significantly higher in the obese and DM groups than in the control group, but there was no significant difference among the obese, TI-DM and TII-DM groups. The study identified a significant relationship between serum SCUBE1 level and hepatic CAS3, IL-6 and SOD expression and pancreatic SOD expression.

**CONCULISON:** SCUBE1 can be used as a promising novel marker for the diagnosis and monitoring of subclinical atherosclerosis in individuals with obesity, TI-DM, and TII-DM

## INTRODUCTION

The global prevalence of diabetes mellitus (DM) and obesity has shown a consistent and sustained rise over the past two decades. Over this period, the prevalence of DM almost doubled, positioning it as the ninth leading cause of mortality ^1–3^. Obesity has emerged as a worldwide pandemic, posing a substantial threat to human lives and constituting a significant public health concern as one of the prevalent non-communicable diseases ^4^. The complications associated with DM have conventionally been categorized into macrovascular complications, such as cardiovascular disease, and microvascular complications, such as complications affecting the kidney, retina and nervous system. Complications in individuals with type II DM (TII-DM) are highly prevalent, with approximately half of TII-DM patients exhibiting microvascular complications and 27% experiencing macrovascular complications ^3^. Among these patients, cardiovascular complications stand as the foremost contributor to morbidity and mortality. Diabetic nephropathy ranks as the primary cause of end-stage renal disease, diabetic retinopathy leads to the most prevalent cases of blindness in developed nations, and diabetic neuropathy represents the predominant risk factor for amputation and the development of foot ulcers ^5^. The critical roles of the endothelium include releasing endothelium-derived vasoactive factors, preventing platelet aggregation and leukocyte adhesion, and regulating cell proliferation. Diabetes-induced endothelial dysfunction is a critical and initiating factor in the genesis of diabetic vascular complications ^6^.

Persistent chronic hyperglycemia and elevated levels of plasma-free fatty acids lead to an increase in oxidative stress (OS) and t proinflammatory cytokines in patients with DM and obesity ^7,8^. Elevations in OS and inflammatory responses cause vascular smooth muscle proliferation, increased collagen formation, and excessive production of growth factors, contributing to accelerated atherosclerosis. ^7,9^. Therefore, many studies have focused on finding an ideal marker that can indicate endothelial dysfunction and chronic inflammation.

Serum signal peptide-CUB-EGF domain-containing protein 1 (SCUBE-1) is a recently characterized cell surface glycoprotein that falls within the superfamily of epidermal growth factors (EGFs). The SCUBE gene family consists of three members, namely SCUBE-1, SCUBE-2, and SCUBE-3, which are categorized as both secretory and membrane proteins ^10^. SCUBE-1 is localized within the alpha granules of endothelial cells and platelets and is released from the cell surface upon thrombin activation. Previously, elevated SCUBE-1 levels have been documented, especially in atherothrombotic conditions ^11–13^ .

The present study conducts a comparative analysis of serum SCUBE1 levels among rats with experimentally induced obesity, type I DM (TI-DM), TII-DM, and a control group. Furthermore, the study includes a histopathological assessment of proinflammatory cytokines such as Caspase-3 (Cas-3), interleukin-6 (IL-6), interferon-gamma (IFN-gamma), and superoxide dismutase (SOD) in the pancreas and liver, and assesses the potential correlations between these inflammatory markers and serum SCUBE. The aim of this study was to investigate whether SCUBE1 can be used as a potential biomarker in the diagnosis and follow-up of subclinical atherosclerosis in obesity and DM.

## MATERIAL AND METHODS

### Animals

Acquired for the study were 28 male Sprague Dawley rats (160-180 g, 4 weeks old) that were housed at a room temperature of 21°C–22°C with 60%± 5% humidity, and in a 12:12-hour light/dark cycle. This experimental study was approved by the Burdur Mehmet Akif Ersoy University Animal Experiments Local Ethics Committee (Ethics number: 17.02.2022/726). Feed (standard rat chow or high-fat diet (HFD), on a case-by-case basis) and water were given ad libitum.

### Experimental procedure

The rats were distributed randomly into four groups, with seven rats in each group, as follows:

1. Control group: The rats were fed standard rat chow (5 weeks) and a single dose of citrate buffer was administered intraperitoneally (i.p.) after overnight fasting for one week before the experiment.
2. HFD group: The rats were fed a HFD (5 weeks) and a single dose of citrate buffer was administered intraperitoneally (i.p.) after overnight fasting for one week before the experiment.
3. HFD+TII-DM group: The rats were fed a HFD (5 weeks) and a single dose of streptozotocin (STZ), 30 mg/kg) dissolved in a 0.01 M citrate buffer (pH 4.5) was administered i.p. after overnight fasting one week before the experiment.

4-TI-DM group: The rats were fed standard rat chow and 25 mg/kg STZ was administered i.p. for 5 consecutive days.

The HFD was prepared locally in our laboratory. The normal rat pellet diet was pulverized and mixed with tallow (1:0.5 ratio), placed in a freezer to solidify and then pelleted.

During the experimental period, the weight gains of the animals were measured every week. Before the start of the study, fasting blood glucose (FBG) measurements were made using a glucometer (Clever Check, Germany) to ensure that all groups were euglycemic, and blood glucose measurements were repeated weekly for the duration of the experiment. At the end of the experiment, rats were decapitated under ketamine (40 mg/kg) and xylazine (5 mg/kg) anesthesia. During necropsy, blood samples were collected from each animal for biochemical analyzes. Serum samples were obtained by centrifugation at 4000 rpm for 10 min and stored at -80°C until analysis, and liver and pancreas samples were collected and fixed in 10% buffered formalin for histopathological and immunohistochemical analysis.

### Histopathological analysis

The fixed liver and pancreas samples were routinely processed using a fully automatic tissue processing device (Leica ASP300S, Leica Microsystems, Wetzlar, Germany) and embedded in paraffin wax. After cooling, 5 μm thick sections were taken from the paraffin blocks using a rotary microtome (Leica RM2155, Leica Microsystems, Wetzlar, Germany) and stained with hematoxylin-eosin (HE), mounted with a coverslip and examined under a light microscope.

### Immunohistochemical analysis

For the immunohistochemical analysis, four series of sections taken from all blocks drawn on poly-L-lysine coated slides were stained immunohistochemically for Cas-3 (Anti-Cas-3 antibody [EPR18297] (ab184787)), IL-6 (anti-IL-6 antibody [1.2-2B11-2G10] (ab9324), INF-gamma (anti-interferon beta antibody (ab140211)) and SOD (anti-SOD 3/EC-SOD antibody [4G11G6] (ab80946)) expression using a streptavidin-biotin peroxidase technique, according to the manufacturer’s instructions. All primary antibodies were used at a 1/100 dilution and purchased from Abcam. The sections were incubated with the primary antibodies for 60 min, and immunohistochemistry was carried out using biotinylated secondary antibodies and streptavidin-alkaline phosphatase conjugates. An EXPOSE Mouse and Rabbit-Specific HRP/DAB Detection IHC kit (ab80436) (Abcam, Cambridge, UK) was used as the secondary antibody. Diaminobenzidine (DAB) was used as the chromogen. For negative controls, an antigen dilution solution was used instead of the primary antibody. All examinations were performed on blinded samples by a specialized pathologist from another university.

For the immunohistochemical analysis, the sections were investigated separately for each antibody. To evaluate the severity of the immunohistochemical reaction of cells with markers, a semiquantitative analysis was performed using a grading score range of (0) to (III) as follows: (0) = negative, (I) = focal weak staining, (II) = diffuse weak staining, and (III) = diffuse marked staining. For evaluation, 10 different areas in each section were examined under 40X objective magnification. Morphometric analyzes and microphotography were performed using the Database Manual Cell Sens Life Science Imaging Software System (Olympus Co., Tokyo, Japan). The results were saved and analyzed statistically.

### Biochemical analysis

All samples were left in an area at room temperature until they were defrosted and then vortexed. Serum glucose, triglyceride, total cholesterol, high-density lipoprotein (HDL) and low-density lipoprotein (LDL) cholesterol, alanine aminotransferase (ALT) and aspartate aminotransferase (AST) levels were determined with an autoanalyzer (Beckman AU5800; Beckman Coulter Diagnostics, Brea, CA).

Serum SCUBE1 levels were quantified using a commercially available ELISA kit (Bioassay Technology Laboratory, Shanghai, China; catalog no. E0948Ra) according to the manufacturer’s instructions. The sensitivity of the assay was 0,35 ng/mL with inter- and intra-assay coefficients of variation of <10 and 8%, respectively. The assay results were expressed in ng/mL

### Statistical Analysis

The between-group comparison of numerical variables was performed using a Kruskal-Wallis test, while post hoc comparisons were made using a Mann-Whitney U-test. The correlation analysis was evaluated with Spearman’s rank test, and the significance of the association between categorical variables was evaluated using a Chi-square test. P values less than 0.05 were accepted as significant.

## RESULTS

### Testing the models of obesity, and STZ-induced TI-DM and TII-DM

The blood glucose levels and weights of the rats included in the study were measured periodically (every 5 weeks). Blood glucose remained stable throughout the follow-up period in the control group, and the weight of the rats in this group increased in parallel with normal development. Blood glucose remained stable in the obese group, while there was an increase in body weight; blood glucose increased together with weight gain in the TII-DM group; and blood glucose increased in the TI-DM group (Figure 1).

**Figure 1:**
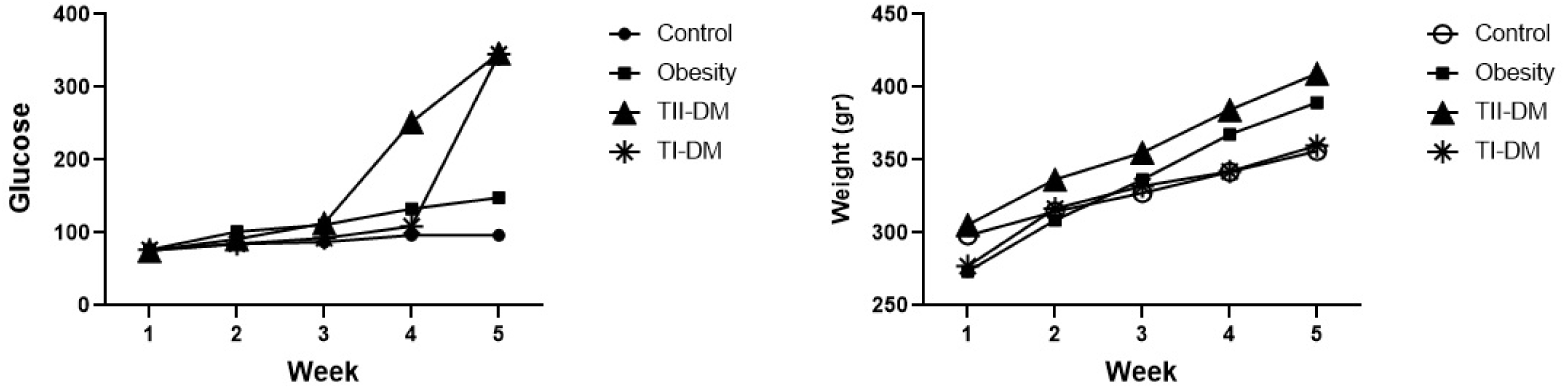
Changes in blood glucose and weight in the experimental groups. Blood glucose increased after week three in the TII-DM group. In the TI-DM group, blood glucose initially showed a slow increase followed by a rapid increase after week four. Body weight increased gradually in the obese and TII-DM groups than in the control and TI-DM groups.

### Biochemical Analyzes

Liver function tests for alanine transaminase (ALT), aspartate transaminase (AST) and albumin were evaluated, as the liver is an important organ in glucose and lipid metabolism. The serum lipid panel included total cholesterol, high-density lipoprotein (HDL) cholesterol, low-density lipoprotein (LDL) cholesterol and triglyceride. The lowest serum albumin levels were observed in the TI-DM group and the highest levels were observed in the TII-DM group (p<0.01). The serum ALT and AST and glucose levels were the highest in the obese group, while the levels were significantly higher in the TI-DM and TII-DM groups than in the control group (p<0.01 for both ALT and AST). As anticipated, the lipid parameters were higher in the obese and TII-DM groups than in the control group and the TI-DM group (p<0.01 for all of the lipid parameters). Serum SCUBE1 levels were higher in the obese, TI-DM and TII-DM groups than in the control group; however, no significant difference was found in the SCUBE1 levels of the obese, TI-DM and TII-DM groups (Table 1). The correlation analysis revealed a positive correlation between serum SCUBE1 levels and AST, ALT, total cholesterol, LDL-cholesterol and triglyceride levels (Table 2).

**Table 1:** Biochemical Parameters.

| PARAMETER | Control (n=7)<br>**(a)** | Obesity (n=7)<br>**(b)** | TI-DM<br>**(c)** | TII-DM<br>**(d)** | **P** |
| --- | --- | --- | --- | --- | --- |
| Albumin<br>(g/L)* | 26.9(25.6-29) | 26.5 (24.2-27.6) | 25 (24.1-26.7) | 31 (27.3-35.2) | **0.004** |
| ALT<br>(U/L)** | 29 (23.1-36.5) | 77.5 (64.5-87.8) | 58 (46.9-67.1) | 33.5 (20.5-71.2) | **<0.01** |
| AST<br>(U/L)*** | 101(88-115.5) | 184 (132.3-185.8) | 124 (106.4-143.5) | 160 (72.2-226.5) | **<0.01** |
| Total<br>Cholesterol<br>(mg/dL)<sup>#</sup> | 39 (33.4-45.1) | 74 (62.7-86.4) | 41 (36.4-51.5) | 93.5(70.5-136.6) | **<0.01** |
| HDL<br>Cholesterol<br>(mg/dL) <sup>##</sup> | 22 (18.4-25.1) | 37.5 (30.7-41) | 25 (21-29.5) | 40.5 (32.9-52.2) | **<0.01** |
| LDL Cholesterol<br>(mg/dL) <sup>###</sup> | 13 (10.5-15) | 30 (22.9-33.4) | 11 (9.4-16.5) | 37 (27.5-65.1) | **<0.01** |
| Triglyceride<br>(mg/dL)<sup>†</sup> | 32 (28.5-63.5) | 106 (91.5-192.4) | 59.5 (48.4-72.5) | 340 (247.2-498.2) | **<0.01** |
| SCUBE1<br>(ng/mL)<sup>††</sup> | 14.6 (6.1-23.5) | 31.8 (28.6-34.7) | 26.6 (15.5-38.7) | 24.6 (18.1-41.2) | **0.01** |
| **post hoc comparisons** |  |  |  |  |  |
| * a>c p=0,04 | d>c p=0,01 | d>c p<0,01 |  |  |  |
| ** b>a p<0,01 | b>c p=0,01 | b>d p=0,01 | c>a p<0,01 | d>a p<0,01 |  |
| *** b>a p<0,01 | b>c p=0,02 | c>a p<0,01 | d>a p<0,01 |  |  |
| <sup>#</sup> b>a p<0,01 | b>c p<0,01 | d>c p<0,01 | d>a p<0,01 |  |  |
| <sup>##</sup> b>a p=0,01 | b>c p=0,01 | d>c p=0,01 | d>a p<0,01 |  |  |
| <sup>###</sup> b>a p=0,04 | b>c p=0,01 | d>c p<0,01 | d>a p=0,01 |  |  |
| <sup>†</sup> b>a p<0,01 | b>c p<0,01 | c>a p=0,04 | d>a p<0,01 | d>b p<0,01 | d>a p<0,01 |
| <sup>††</sup> b>a p<0,01 | c>a p=0,04 | d>a p=0,02 |  |  |  |

ALT: Alanine aminotransferase , AST: Aspartate aminotransferase , HDL High-density lipoprotein, LDL Low-density lipoprotein, SCUBE1: Signal Peptide-CUB Domain-EGF-Related 1 Protein

**Table 2:**
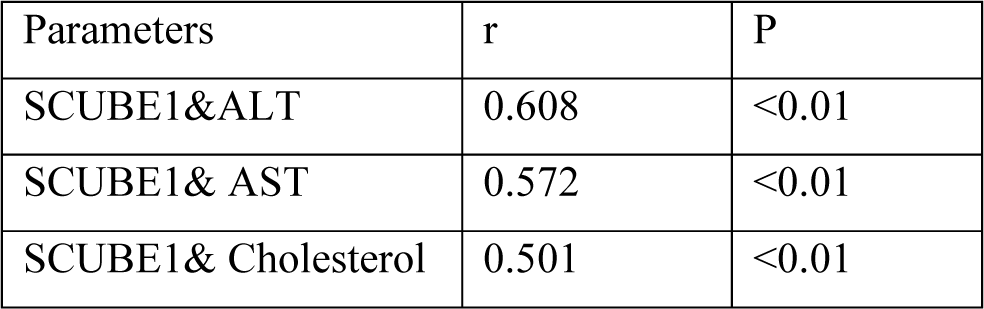
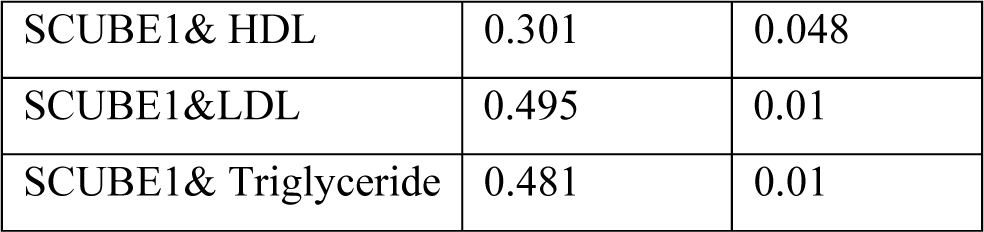
The correlations between biochemical parameters.

| Parameters | r | P |
| --- | --- | --- |
| SCUBE1&ALT | 0.608 | <0.01 |
| SCUBE1& AST | 0.572 | <0.01 |
| SCUBE1& Cholesterol | 0.501 | <0.01 |
| SCUBE1& HDL | 0.301 | 0.048 |
| SCUBE1&LDL | 0.495 | 0.01 |
| SCUBE1& Triglyceride | 0.481 | 0.01 |

### Histopathological analysis

The histopathological examination revealed completely normal liver and pancreas tissue in the control group, and apparent hyperemia and degeneration in the obese, TI-DM and TII-DM groups. Moderate-to-severe lipidosis was observed in the hepatocytes of the obese rats fed a HFD, and similarly, increased vacuolar degeneration was noted in the obese, TI-DM and TII-DM groups (Figure 2).

**Figure 2:**
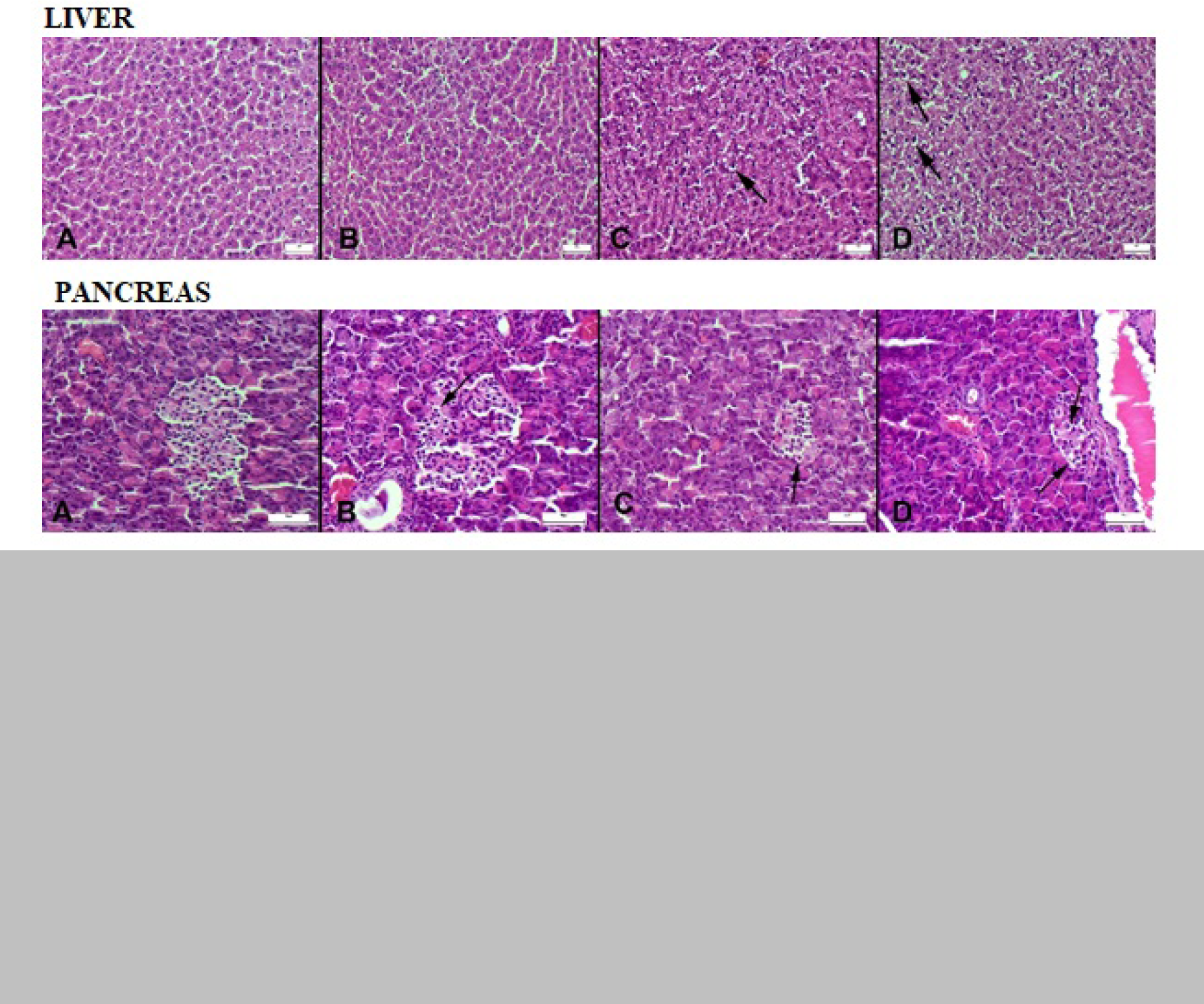
Histological changes in the liver and pancreas. **Liver:** (A) Normal liver histology in the control group. (B) Slight degenerative changes in the TI-DM group. (C) Moderate lipidosis (arrow) in the obesity group. (D) Severe degeneration and lipidosis (arrows) in TII-DM. **Pancreas:** (A) Normal pancreatic histology in the control group. (B) Slight vacuolar degenerative changes in cells (arrow) in the TI-DM group. (C) Increased degenerative changes in the cells (arrow) in the Obesity group. (D) Marked increase in degenerative changes (arrows) in cells in the TII-DM group. HE, scale bar=50µm.

### Immunohistochemical Findings

The expression of CAS-3, as a marker of inflammation, the expression of the inflammatory markers IL-6 and INF-gamma, and the expression of SOD, a marker of OS, were evaluated in an immunohistochemical examination of liver and pancreas tissues. SCUBE1 immunoexpression was not evaluated in the liver or pancreas parenchyma, as these organs lack SCUBE1 expression.

In the immunohistochemical examination of the liver, Cas-3, IL-6 and INF-gamma expression was increased in the obese, TI-DM and TII-DM groups, while there was mild or no expression in the control group. SOD expression was decreased in the obese, TI-DM and TII-DM groups compared with in the control group (Figure 3).

**Figure 3:**
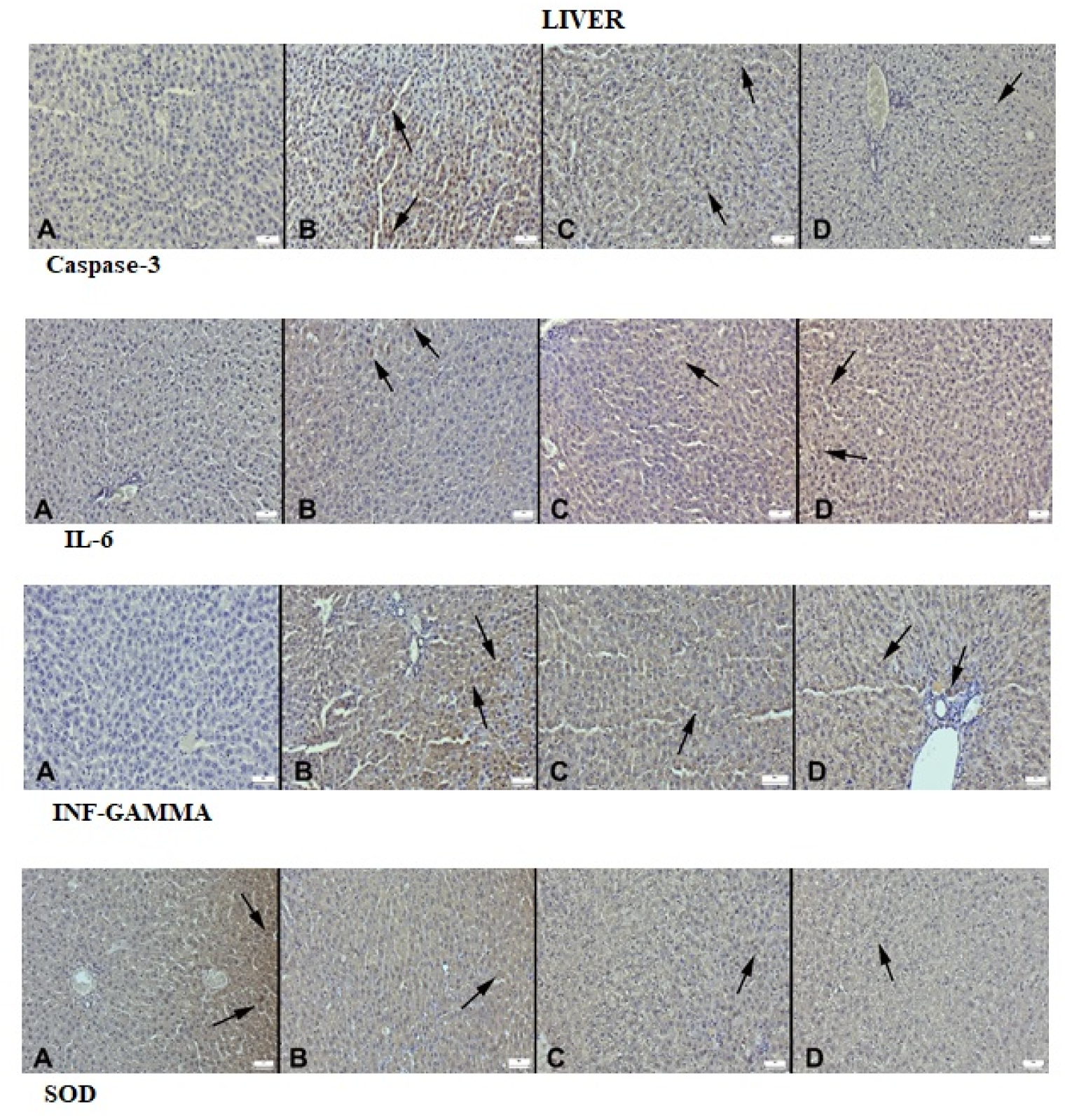
Immunohistochemical marker expressions in the liver. **Caspase 3 expressions between groups** (A) Negative expression in the control group. (B) Increased expression (arrows) in the TI-DM group. (C) Moderate expression (arrows) in the obesity group. (D) Slight expression (arrow) in the TII-DM group. **IL-6 expressions between groups**: (A) Negative expression in the control group. (B) Increased expression (arrows) in the TI-DM group. (C) Moderate expression (arrow) in the obesity group. (D) Increased expression (arrows) in the TII-DM group. **INF-gamma expressions among the groups:** (A) No expression in the control group. (B) Increased expression (arrows) in the TI-DM group. (C) Moderate expression (arrow) in the obesity group. (D) Marked increased expression (arrows) in the TII-DM group. **SOD expressions among the groups**: (A) Marked expression in the control group. (B) Decreased expression (arrow) in the TI-DM group. (C) Moderately decreased expression (arrow) in the obesity group. (D) Marked decreased expression (arrow) in the TII-DM group (Streptavidin biotin peroxidase method, scale bars=50µm).

Similarly, an immunohistochemical examination of pancreas tissue showed increased Cas-3, IL-6 and INF-gamma expressions in the obese, TI-DM and TII-DM groups, while there was zero, or only mildly increased expression in the control group. SOD expression was decreased in the obese, TI-DM and TII-DM groups compared with control group (Figure 4).

**Figure 4:**
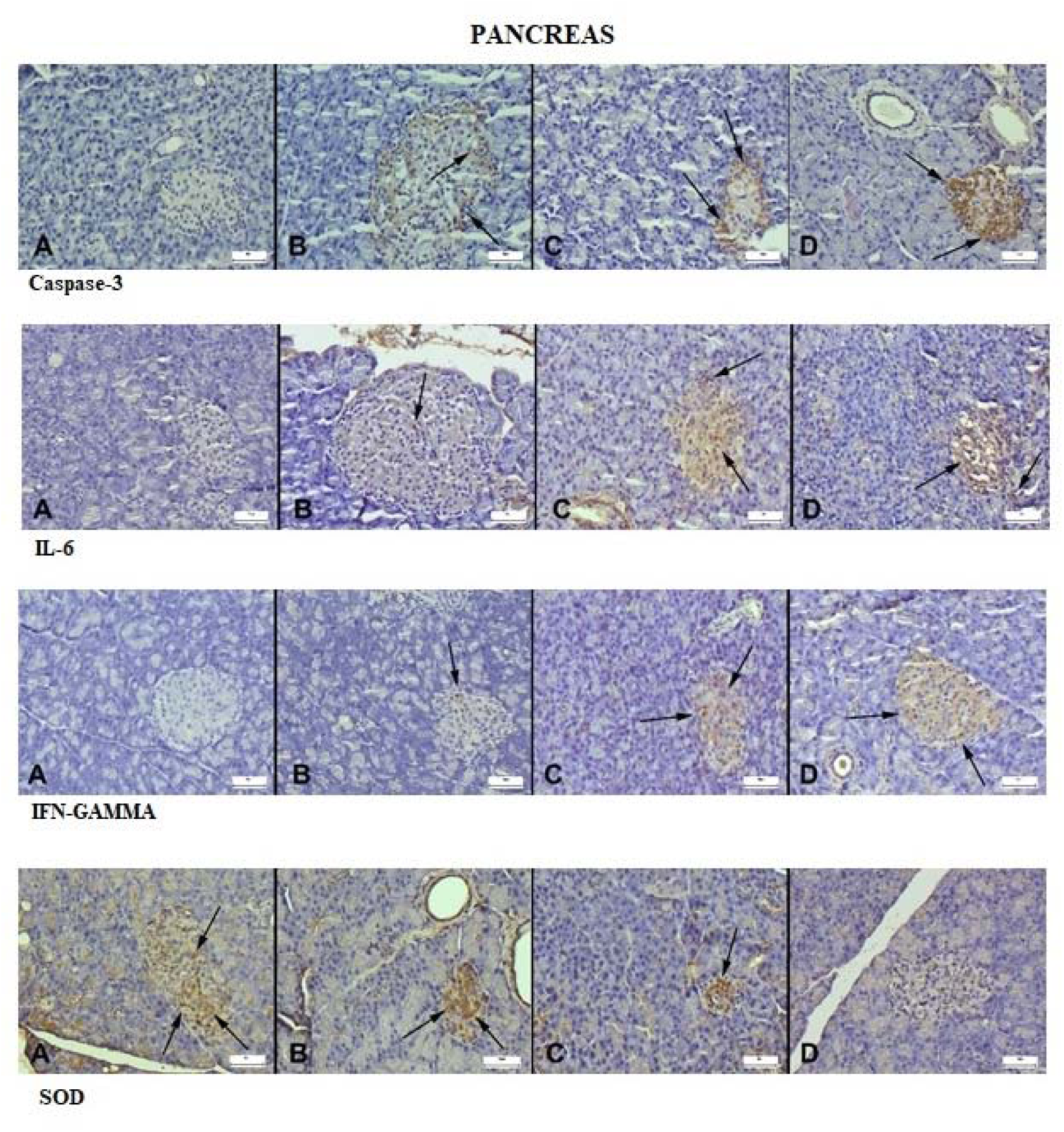
Immunohistochemical marker expressions in the pancreas. **Caspase-3 expressions in the pancreas between the groups** (A) Negative expression in the control group. (B) Increased expression (arrows) in the TI-DM group. (C) Moderate expression (arrows) in the obesity group. (D) Marked increase in expression (arrow) in the TII-DM group. **IL-6 expressions in pancreases between groups**: (A) No expression in the control group. (B) Slight expression (arrow)the in TI-DM group. (C) Increased expression (arrows) in the obesity group. (D) Marked increase in expression (arrows) in the TII-DM group. **Pancreatic INF-gamma expressions between groups:** (A) Negative expression in the control group. (B) Slight increase in expression (arrow) in the TI-DM group. (C) Moderate increase in expression (arrows) in the obesity group. (D) Marked increase in expression (arrows) in the TII-DM group. **SOD expressions in pancreas among the groups**: (A) Marked expression (arrows) in the control group. (B) Decreased expression (arrows) in the TI-DM group. (C) Moderately decreased expression (arrow) in the obesity group. (D) Almost negative expression in the TII-DM0 group (Streptavidin biotin peroxidase method, scale bars=50µm).

A semi-quantitative analysis was carried out to evaluate the intensity of immunohistochemical reactions on a scale of 0–III: 0=negative, I=focal weak staining, II=diffuse weak staining and III=diffuse strong staining. Cas-3, IL-6 and INF-gamma expression in the liver was noted to increase, while SOD expression decreased in the obese, TI-DM and TII-DM groups than in the control group (p<0.001 for all). Similarly, Cas-3, IL-6 and INF-gamma expression in pancreatic tissue were increased while SOD expression was decreased in the obese, TI-DM and TII-DM groups compared with control group (p= 0.004 for IL-6, p<0.001 for the others). (Table 3)

**Table 3:** Statistical analysis results of the liver and pancreas immunohistochemical scores.

|  |  | Control |  |  |  | Obesity |  |  |  | TI-DM |  |  |  | TII-DM |  |  |  | p |
| --- | --- | --- | --- | --- | --- | --- | --- | --- | --- | --- | --- | --- | --- | --- | --- | --- | --- | --- |
|  |  | 0 | I | II | III | 0 | I | II | III | 0 | I | II | III | 0 | I | II | III |  |
| **LIVER** | Cas-3 (n) | 7 | 1 | 0 | 0 | 0 | 5 | 3 | 0 | 0 | 4 | 4 | 0 | 0 | 6 | 2 | 0 | <0.001 |
|  | IL-6 (n) | 7 | 1 | 0 | 0 | 0 | 4 | 4 | 0 | 0 | 5 | 3 | 0 | 0 | 2 | 6 | 0 | <0.001 |
| | INF-γ (n) | 7 | 1 | 0 | 0 | 0 | 0 | 6 | 2 | 0 | 2 | 6 | 0 | 0 | 0 | 4 | 4 | <0.001 |
|  | SOD (n) | 0 | 3 | 5 | 0 | 2 | 5 | 1 | 0 | 0 | 6 | 2 | 0 | 5 | 3 | 0 | 0 | <0.001 |
| **PANCREAS** | Cas-3 (n) | 8 | 0 | 0 | 0 | 0 | 7 | 1 | 0 | 5 | 3 | 0 | 0 | 0 | 1 | 5 | 2 | <0.001 |
|  | IL-6 (n) | 7 | 1 | 0 | 0 | 2 | 6 | 0 | 0 | 5 | 3 | 0 | 0 | 0 | 4 | 4 | 0 | 0.004 |
| | INF-γ (n) | 8 | 0 | 0 | 0 | 0 | 5 | 3 | 0 | 4 | 4 | 0 | 0 | 0 | 1 | 7 | 0 | <0.001 |
|  | SOD (n) | 0 | 0 | 6 | 2 | 2 | 6 | 0 | 0 | 0 | 3 | 5 | 0 | 6 | 2 | 0 | 0 | <0.001 |

For the immunohistochemical analysis, sections were investigated separately for each antibody. To evaluate the severity of the immunohistochemical reaction of cells with markers, a semiquantitative analysis was performed using a grading score of (0)–(III) as follows: (0) = negative, (I) = focal weak staining, (II) = diffuse weak staining, (III) = diffuse strong staining. Cas-3: Caspase 3, IL-6: Interleukin-6, INF-γ : interferon gamma, SOD: superoxide dismutase. Chi square analysis.

The study also included a semi-quantitative analysis of Cas-3, IL-6, INF-γ and SOD expression in an immunohistochemical examination of liver and pancreatic tissues to identify any relationship with serum SCUBE1 levels. Serum SCUBE1 levels were found to increase in parallel to increasing Cas-3 and IL-6 expression in the liver, and inversely, SCUBE1 levels were found to decrease with increasing SOD expression. Although SCUBE1 levels increased with increasing INF-gamma expression in the liver, the increase was not statistically significant, and although statistically insignificant, the Cas-3, IL-6 and INF-gamma expressions in the pancreatic tissue increased with increasing serum SCUBE1 levels. A significant decrease in pancreatic SOD expression was also noted with increasing serum SCUBE1 levels (Figure-5 post hoc analysis).

**Figure 5:**
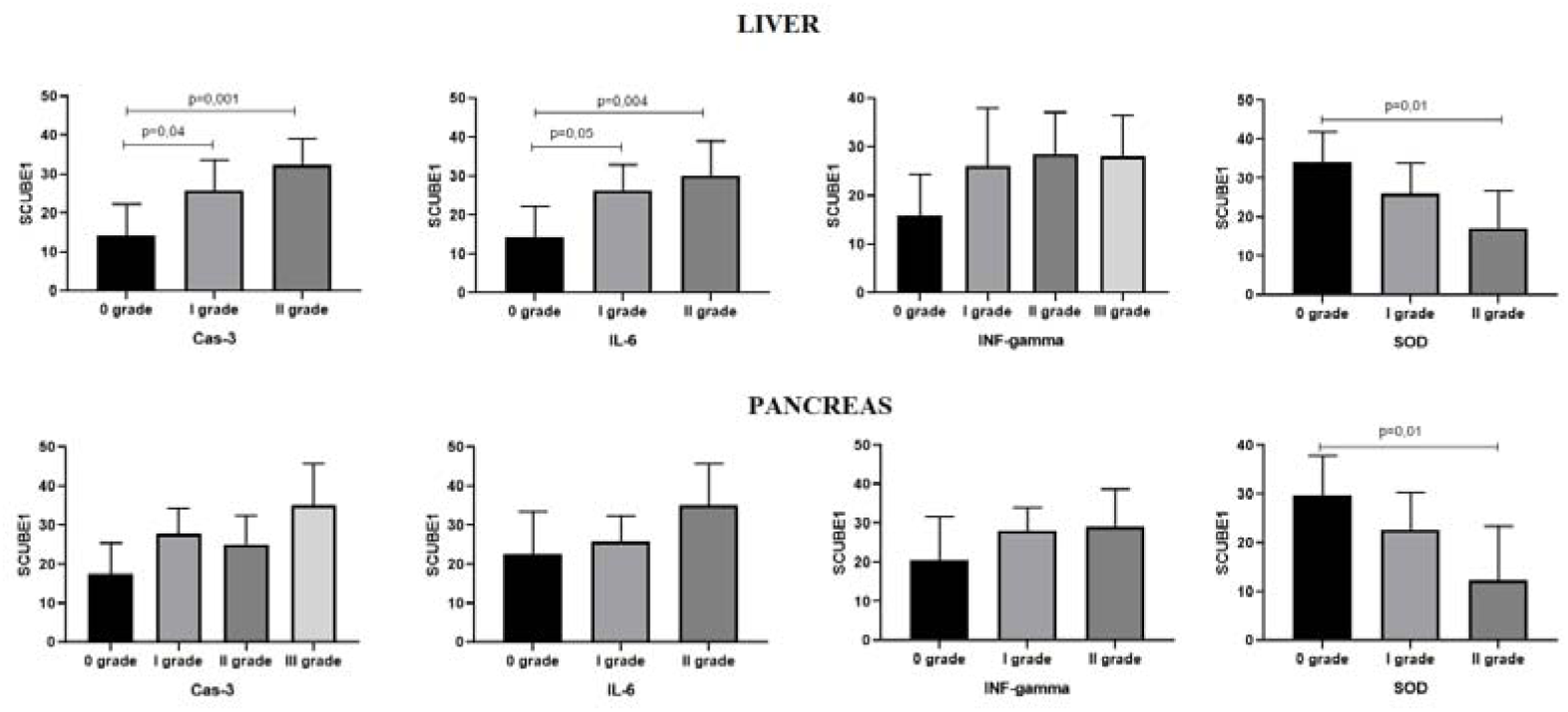
Relationship between liver and pancreas immunohistochemical markers and serum SCUBE1 levels. **In the liver**, for Cas-3: serum SCUBE1 levels are grade II> grade 0 (p=0.001) and grade I> grade 0 (p=0.04). For IL-6: serum SCUBE1 levels are grade II> grade 0 (p=0.004) and grade I> grade 0 (p=0.05, near significant). For INF-gamma Serum SCUBE1 levels are lowest at grade 0, but with no statistical significance. For SOD, serum SCUBE1 levels are grade 0> grade II (p=0.01). **In the pancreas**, serum SCUBE1 levels are lowest at grade 0 for Cas-3, IL-6 and INF-gamma, but with no statistical significance. Only for SOD, serum SCUBE1 levels are grade 0> grade II (p=0,01).

## DISCUSSION

Considering that obesity is the most widespread non-communicable disease worldwide and that one in every 11 people globally has DM, it becomes evident how crucial it is to promptly identify complications arising from these diseases ^4^. To fulfill this objective, we examined the potential of SCUBE-1, which we deem as a highly suitable candidate for a biomarker. We observed that serum SCUBE1 levels were significantly elevated in the obese and DM groups compared to the control group. However, there was no statistically significant difference in SCUBE1 levels among these groups. In histopathological analyzes, various degrees of degenerative changes were identified in the liver and pancreas of the obese and DM groups compared to the control group. Immunohistochemical examinations indicated an increase in Cas-3 and IL6 expression, along with a decrease in IFN-gamma and SOD expression, in the liver and pancreatic tissues of the obese and DM groups. In addition, we detected a significant correlation between serum SCUBE1 levels and hepatic CAS3, IL-6, and SOD expression as well as pancreatic SOD expression.

SCUBE1, which plays an important role in embryonic development and differentiation, is expressed mostly in the endothelium and platelets in adult tissues ^11,14,15^. As SCUBE1 is strongly localized in the platelet- and fibrin-rich areas in the organized thrombus, it has been suggested to be a strong marker of thrombus formation, and it has also been well documented that activated platelets play an important role in the onset and progression of atherosclerosis ^16–19^. Subsequent clinical studies have revealed elevated SCUBE1 levels in the sera of patients with acute coronary syndrome and acute stroke, and serum SCUBE1 has been identified as an independent risk factor for acute coronary syndrome ^20–22^. One or more single-nucleotide polymorphisms have been identified in the SCUBE1 gene of patients with venous thromboembolism, and studies have also reported elevated serum SCUBE1 levels in patients with pulmonary embolism ^23,24^. Furthermore, SCUBE-1 levels were decreased by treatments against IL-1β and TNF-α (11).

Endothelial dysfunction, which represents a pivotal cardiovascular risk factor, has been the subject of comprehensive investigation in both diabetic animals and human subjects. This is due to its significant role as a primary and initiating factor in the development of vascular diseases. Due to SCUBE-1 is stored in alpha granules of the endothelium, it is considered a potential indicator for evaluating endothelial dysfunction. Research studies conducted with this premise have also provided confirmation. For instance, in a study by Tüfekci et al. SCUBE1 was identified as a viable marker for detecting preclinical endothelial dysfunction in patients with acromegaly ^25^. However, there are only a few and conflicting outcome studies investigating the connection between SCUBE1 and vascular complications arising from obesity and DM in the literature. For instance, Bingol et al. conducted a comparative analysis involving type 2 diabetic patients with or without complications, as well as healthy individuals. Their findings revealed that SCUBE-1 levels exhibited no statistically significant variance among the groups. However, a positive correlation was observed between serum SCUBE-1 levels and fasting blood glucose levels. It is noteworthy that, although not reaching statistical significance, the study by Bingol et al. noted the highest SCUBE-1 levels among diabetic patients with microvascular complications ^26^. In addition Icel et al. reported elevated SCUBE1 levels in patients with TII-DM ^27^. Bodur and et al. examined the association between OS and SCUBE1 levels in in obese rats induced by a HFD. They observed that SCUBE1 levels were lower in obese rats than in the control group. In addiditon they found that serum triglyceride levels increased but glucose levels decreased in the obese group and both triglyceride and glucose levels exhibited no significant difference between the control and obese groups ^28^. Dai et al. observed that plasma SCUBE1 levels were higher in cases of acute coronary syndrome and acute ischemic stroke compared to the control group, whereas Şahin et al. did not identify a statistically significant correlation ^29,30^. In the current study, we found that SCUBE 1, triglyceride and glucose levels were higher in obese and DM rats. The serum SCUBE1 level can be influenced by various factors, including blood pressure, medication usage, hypoxia, and physical activity. These variables were not taken into account sufficiently in the studies conducted by Bingol et al. and Icel et al. ^26,27^. In our study, it was quite homogenized in this respect. Metabolic parameters can fluctuate based on the specific breed, gender, and duration of dietary intake in the animal model ^31^. In the study by Bodur et al., they noticed that glucose levels were lower in obese rats, despite the fact that these rats were exposed to a HFD for twice the duration compared to our study (10 weeks versus 5 weeks) ^28^. This dietary regimen and the resulting low glucose levels may be responsible for the low detection of SCUBE1.

As mentioned earlier, despite its original discovery in the endothelium, SCUBE1 is also highly expressed in platelets. Previous studies have shown SCUBE1 to be mostly localized in the alpha granules and the open canalicular system of platelets, and it has been shown that a certain amount of SCUBE1 attaches to the membrane while a large amount is retained in the alpha granules of platelets ^16^. In vitro studies have shown that the membrane-bound form of SCUBE1 stored in the alpha granules of platelets is exposed to the platelet surface upon platelet activation (i.e. CD40L and P-selection) ^16^. For instance, Wu et al. observed that in mice lacking plasma SCUBE1 (Δ/Δ mice), standard hematologic and coagulation characteristics, as well as the expression of major platelet receptors, remained unaltered. However, Δ/Δ platelet-rich plasma exhibited diminished platelet aggregation when exposed to ADP and collagen. Interestingly, the introduction of purified recombinant SCUBE1 protein restored platelet aggregation in Δ/Δ platelet-rich plasma. A deficiency of SCUBE1 in plasma reduced arterial thrombosis occurrence in mice and provided protection against fatal thromboembolism initiated by collagen-epinephrine treatment. Additionally, antibodies targeting the epidermal growth factor–like repeats of SCUBE1, which participate in trans-homophilic protein-protein interactions, safeguarded mice from lethal thromboembolism ^32^. These studies ultimately demonstrated that soluble plasma SCUBE1 is not only a marker but also an active participant in thrombosis.

Previous studies have identified an increase in soluble P-selectin and CD40 ligands — found together with SCUBE1 — in the alpha granules of platelets in cases of acute hyperglycemia ^33,34^. The direct osmotic effect of hyperglycemia can also play a role in platelet activation, and studies have identified increased platelet protein C kinase activity, which is pro-aggregatory both in chronic and acute hyperglycemia ^35,36^. Furthermore, nonenzymatic glycation can occur as a result of hyperglycemia in platelet membrane proteins, and as a result, platelet plasma membrane fluidity is decreased and sensitivity to platelet aggregation agonists is increased in diabetic patients ^37,38^. Increased platelet activation has also been observed in obese patients ^39^. Various markers of platelet activation are increased in obese patients and those with DM, in whom markers of platelet hyperactivation, including platelet microparticle levels in the circulation, thromboxane B2 metabolites, soluble P-selectin and platelet-derived CD40L, have been found to be increased. Platelet-dependent thrombosis is increased by leptin, the hormone essentially synthesized by the adipose tissue, leading to the sensation of satiety, and is decreased by adiponectin, an insulin-sensitive adipokine produced solely by adipocytes. It has been reported that insulin receptor-mediated signaling sensitizes platelets against the aggregation-preventing effects of prostaglandin I2 and nitric oxide ^40^. Furthermore, tissue factor expression, which is dependent on platelet adhesion and thromboxane A2, is inhibited by insulin in healthy patients, and is increased in obese patients with insulin resistance ^40^. Lipid metabolism can also alter platelet function. It has been demonstrated that hypertriglyceridemia and the resulting high very low-density lipoprotein (VLDL) levels and the associated high apolipoprotein E levels play an important role in platelet activation through the LDL receptor found on the platelets ^41^. The present study identified high serum SCUBE1 levels in obese,DM rat models. SCUBE1 may be considered a new marker of platelet dysfunction and the associated pathological conditions occurring in patients with obesity, TI-DM and TII-DM. As a result, there is a tendency for increased platelet aggregation in obese and diabetic patients. Consequently, we hypothesized that SCUBE1 could be an ideal marker for detecting subclinical atherosclerosis. In fact, our study revealed higher serum SCUBE1 levels in obese and diabetic patients

As mentioned above, microvascular complications primarily exhibit endothelial dysfunction, which is marked by a decrease in nitric oxide (NO) release, increased OS, elevated production of inflammatory factors, abnormal angiogenesis, and impaired endothelial repair. Reactive oxygen species (ROS) and reactive nitrogen species (RNS) are generated as part of normal physiological processes. Nevertheless, the production of ROS escalates under specific pathophysiological circumstances, including obesity, hyperglycemia, and type II diabetes. The heightened production of ROS, coupled with insufficient antioxidant enzyme systems such as catalase, glutathione peroxidase (GPx), and SOD, typically culminates in an imbalance between oxidants and antioxidants, ultimately resulting in OS (27). Elevated OS levels and intensified inflammatory reactions are collaborators in inducing vascular dysfunction associated with diabetes, particularly concerning microvascular complications. Serum concentrations of cytokines and inflammatory markers (such as tumor necrosis factor [TNF]-α, interleukin [IL]-6, soluble CD40L, soluble ICAM, and high-sensitivity C-reactive protein) are elevated in both diabetic animals and human subjects (7).

Consistent with the prior literature, our study revealed increased Cas-3, IL-6, and INF-gamma expression in the liver and pancreatic tissues and decreased SOD expression in the obese and DM groups. Furthermore, we established a noteworthy correlation between serum SCUBE1 levels and the expression of hepatic Cas-3, IL-6, and SOD, as well as pancreatic SOD, suggesting a relationship between serum SCUBE1 levels and the inflammatory process in the liver and pancreas. Finally, serum ALT AST glucose and cholesterol (total cholesterol, LDL-cholesterol and triglyceride) levels were higher in the obese and DM groups than in the control group. A positive correlation was also detected between serum SCUBE1 levels and these laboratory parameters.

The present study has some limitations, such as the effect of the STZ used to induce a model on endothelial cells, liver and other organs (while no STZ was used in the obesity model), and the fact that the expression of SCUBE1 and other inflammatory markers in the rat vessel wall and the platelet function of the rats were not examined. Considering the limitations of the present study, there is a need for further experimental and large-scale studies involving patients with obesity, TI-DM and TII-DM.

In conclusion, this study demonstrated an increase in SCUBE1 levels in rats subjected to experimental induction of obesity, TI-DM), and TII-DM. This elevation was found to be correlated with an increase in inflammatory markers and a decrease in antioxidants within the liver and pancreas. Consequently, our findings suggest that SCUBE1 can be used as a valuable marker for assessing endothelial dysfunction and platelet hyperactivity which are the most crucial initiating factors of obesity and DM-related complications, in other words, in the diagnosis and follow-up of subclinical atherosclerosis.

### Affiliations

Department of Internal Medicine, Antalya Training and Research Hospital of Ministry of Health, Turkey (M.K.). Department of Physiology, Faculty of Medicine, Suleyman Demirel University, Isparta, Turkey (R.A.). Department of Pathology, Faculty of Veterinary Medicine, Mehmet Akif Ersoy University, Burdur, Turkey (Ö.Ö.). Department of Medical Pharmacology, Faculty of Medicine, Akdeniz University, Antalya, Turkey (D.D.). Department of Clinical Biochemistry, Antalya Training and Research Hospital of Ministry of Health, Antalya, Turkey (H.Y.E.).

## Acknowledgements

Not applicable

## Author contribution statement

Study Design: HYE, MK, RA Conceptualization :HYE, MK Methodology and data collection RA and ÖÖ Data interpretation and curation: HYE, MK RA ÖÖ and DD. Formal analysis HYE and MK. HYE, MK, RA,ÖÖ and DD Original draft, and writing: HYE,MK,RA Writing: Review and Editing: All authors.

All authors had full access to all the data in the study and all had final responsibility for the decision to submit for publication. The authors vouch for the accuracy and completeness of the data and for the adherence of the trial to the protocol

## NONSTANDARD ABBREVİATİONS AND ACRONYMS

Cas-3: caspase-3
DM: diabetes mellitus
HFD: high-fat diet
INF-gamma: interferon-gamma
IL-6: interleukin-6
OS: oxidative stress
SCUBE-1: serum signal peptide-CUB-EGF domain-containing protein 1
SOD: superoxide dismutase
STZ: streptozocin

